# SubFeat: Feature Subspacing Ensemble Classifier for Function Prediction of DNA, RNA and Protein Sequences

**DOI:** 10.1101/2020.08.04.228536

**Authors:** H.M.Fazlul Haque, Fariha Arifin, Sheikh Adilina, Muhammod Rafsanjani, Swakkhar Shatabda

**Affiliations:** Department of Computer Science and Engineering, United International University, Plot-2, United City, Madani Avenue, Badda, Dhaka-1212, Bangladesh

**Keywords:** Feature Subspacing, Ensemble Classifier, Biological Entities, Machine Learning, Classification

## Abstract

The information of a cell is primarily contained in Deoxyribonucleic Acid (DNA). There is a flow of information of DNA to protein sequences via Ribonucleic acids (RNA) through transcription and translation. These entities are vital for the genetic process. Recent developments in epigenetic also show the importance of the genetic material and knowledge of their attributes and functions. However, the growth in known attributes or functionalities of these entities are still in slow progression due to the time consuming and expensive *in vitro* experimental methods. In this paper, we have proposed an ensemble classification algorithm called *SubFeat* to predict the functionalities of biological entities from different types of datasets. Our model uses a feature subspace based novel ensemble method. It divides the feature space into sub-spaces which are then passed to learn individual classifier models and the ensemble is built on this base classifiers that uses a weighted majority voting mechanism. *SubFeat* tested on four datasets comprising two DNA, one RNA and one protein dataset and it outperformed all the existing single classifiers and as well as the ensemble classifiers. *SubFeat* is made availalbe as a Python-based tool. We have made the package *SubFeat* available online along with a user manual. It is freely accessible from here: https://github.com/fazlulhaquejony/SubFeat.

## 1 Introduction

With the advent of modern sequencing machines and techniques there had been a tremendous growth in the know sequences. DNA, RNA and proteins are of primary interest. They are involved in all information flow and even in epigenetics. A huge number of sequences and their attributes and properties are very vital to understand the cell organisms. Among these are structure [1], gene-coding markers [2, 3], anti-cancer properties [4], editing [5], binding[6, 7], post-translational modifications [8, 9, 10], sub-cellular localization [11], methylation [12], and many other important process and functions that regulates almost all the processes within the cell organism. However, these techniques are time consuming and expensive.

There have been growth in developing computational and knowledge based methods to predict the attributes and functions of the sequences [13, 14, 15, 16, 17]. One of the key advantages of the knowledge based methods is that they often provide further insights to the patterns that are discoverable using fast computational facilities available and even with relatively small amount of data knowledge transfers and deep learning are also been possible from one problem to another [18, 19, 2, 20]. One of the common approaches in the literature is to formulate the prediction task as a supervised learning problem: binary [21] or multi-class [2] or multi-label [22]. A number of successful classifiers have been used, single classifiers like Support Vector Machines (SVM) [22], K-Nearest Neighbors (KNN) [23], Decision Trees (DT) [24], Naive Bayes (NB) [25], Logistic Regression (LR) [26] and ensemble methods like AdaBoost [27], Random Forest [28], etc have been applied to solve these problems. However, no single method seems to be performing well over other mehods, there are scope to develop new techniques.

One of the most important factor in building a successful machine learning based method is the representation of the dataset. In this case, its how the sequences of DNA, RNA and proteins are converted to vector representation. Generally, ensemble methods are found to provide superior performances provided that they utlize the underlying feature space properly. AdaBoost iteratively learns using weak classifiers, however the algorithm does not exploit or consider the underlying feature space. On the other hand Random Forest smaples the features in a randomly way. From the point of view of biological domain, it has been often seen that in many cases, the features are grouped into several sub-groups based on their respective generating techniques and sometimes the subgroups too share important knowledge. Our main idea in this work is to utilize this property of the feature space.

In this paper, we present a ensemble method called *SubFeat*. *SubFeat* divides the full feature space into overlapping or non-overlapping sub-spaces and learns base classifiers or their mix on the subspaces and the ensemble is created using a voting techniques. It is much similar to Random Forest or Ensemble Voting technique in the way how it uses the feature space and the voting mechanism. However, the approach taken to divide the subspace is unique here. We have tested the problem to four problems related to DNA, RNA and proteins: DNA-binding proteins prediction using protein sequences, A-to-I editing prediction of RNA sequences and promoter and recombination hotspot prediction of DNA sequences. The datasets used in the work are all standard benchmark datasets. The feature space or feature representation used here is generated solely from sequences. The experimental results shows the superiority of the proposed method, *SubFeat* over several single classifiers and ensembles. We have made the methodology available as a Python package freely available and usable from: https://github.com/fazlulhaquejony/SubFeat.

## 2 Materials and Methods

The basic idea of the ensemble method, *SubFeat* is given in Figure 1. In this paper, we have divided the feature space intro three sub-spaces. Each were then trained using a base classifier and the final prediction is made based on the weighted majority voting of the sub-classifiers. The framework is capable of utilizing the possible overlap or non-overlap among the feature spaces.

**Figure 1:**
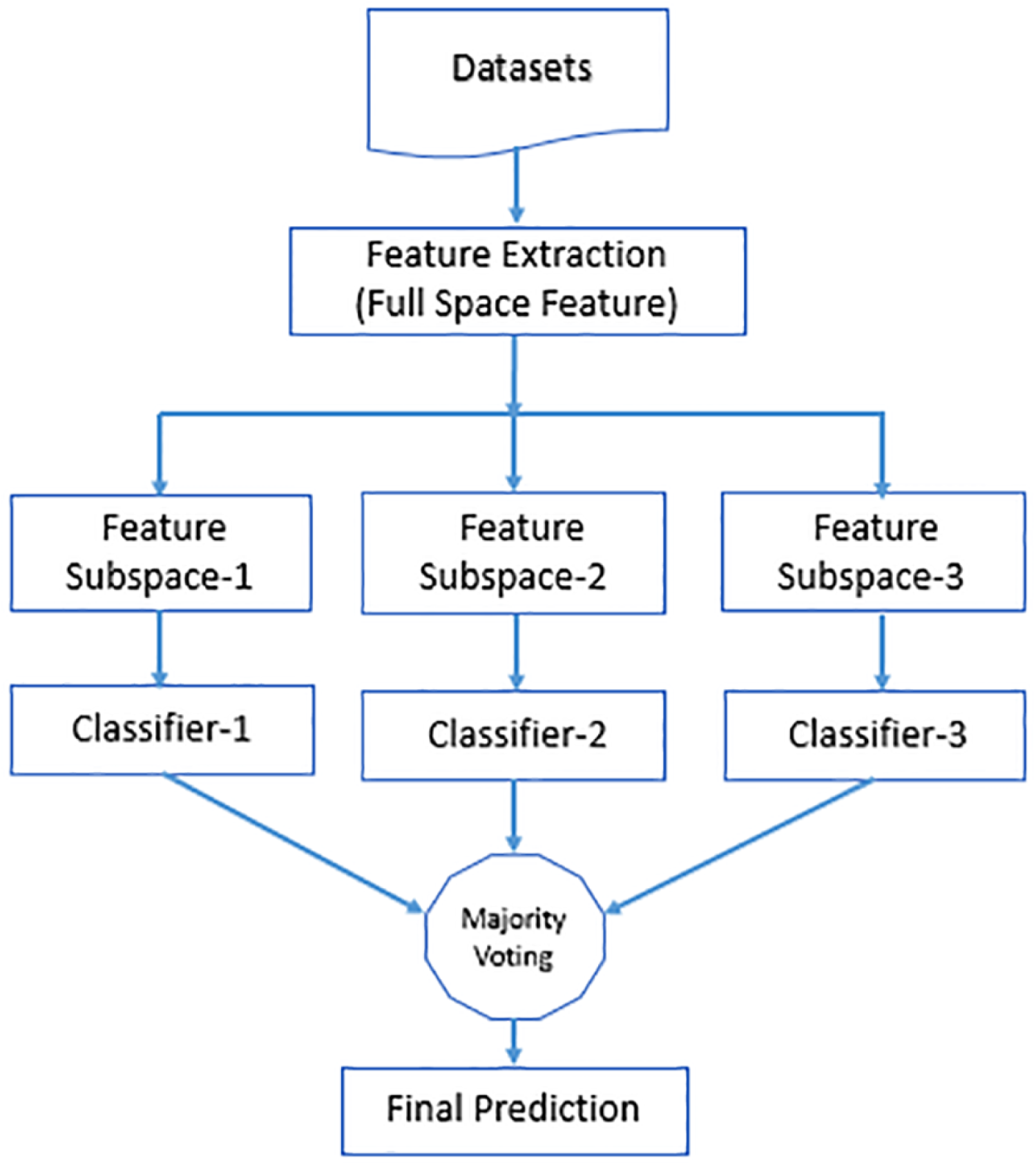
Block diagram for ensemble classifier

In this section, we provide the details of our methods and materials. The section starts with a description of the datasets and the problems that were selected for experiments. A very brief literature review from the computational point of view is also provided for each problem. After that, we describe our feature representation for each of the problems. The ensemble is presented next with the choice of the algorithms in brief. We also describe the performance evaluation techniques used for the work.

### 2.1 Datasets

For this work, we have considered four problems: prediction of DNA recombination hotspots, predition of promoter sequences in DNA, RNA A-to-I editing prediction and prediction of DNA binding proteins. Thus we have incorporated three types of the sequences: DNA, RNA and proteins. In this section, we provide a description of the dataset collection and a brief literature review of the state-of-the-art methods of each of the problems. In supervised machine learning a dataset is generally composed of positive and negative samples.

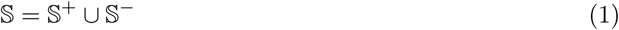

Here, 𝕊^+^ denotes the set of positive instances and 𝕊^−^ denotes the set of negative examples. In this work, we have selected only balanced set of examples where none of the classes positive or negative outnumber the other. A summary of the datasets used in this paper is given in Table 1.

**Table 1:** Summary of the different datasets used to test the performance of *SubFeat*.

| Dataset | Sequence Type | Positive Instances | Negative Instances | Total |
| --- | --- | --- | --- | --- |
| Recombination Hotspot | DNA | 478 | 572 | 1050 |
| $\sigma^{70}$ promoters | DNA | 741 | 1400 | 2141 |
| RNA Editing | RNA | 125 | 119 | 244 |
| DNA Binding Proteins | Protein | 525 | 550 | 1075 |

#### 2.1.1 Recombination Hotspot

Hotspots are regions in the genome where rates of meiotic recombination is much higher compared to the cold spots. DNA binding arrays are used *in vitro* to find recombination hot spots [29]. The dataset that we consider in this paper was originally curated by Jiang et al. [30]. Recently, a good number of machine learning based algorithms and methods [31, 32] as well as ensemble based methods [33] are being proposed in the literature to solve the problem computationally. In this dataset, there were 478 positive samples and 572 negative samples after removing redundancy using CD-HIT [34].

#### 2.1.2 *σ*^70^ Promoters

Promoters are regions in the DNA where RNA polymerase binds itself initiating the transcription process. The RNA polymerase combines itself with different *σ* factors which are differentiated according to their nuclear weights. *σ*^70^ factors are primary house keeping factors and hence have potential importance in gene transcription. The dataset that we have selected here for promoter sequence prediction is taken from [35]. Originally there were curated from RegulonDB [36]. In recent years, we a large number of methods have been proposed to solve the promoter detection problem using this dataset [35, 37, 3, 38]. In this dataset, the promoter sequences are all DNA short sequences and there are 741 positive and 1400 negative sequences.

#### 2.1.3 RNA Editing

Adenosine to Inosine (A-to-I) editing is one of the most common and important RNA modiffications [39] that changes the gene templates and thus affects the genetic variation in species. RNA-DNA difference (RDD) methods are generally employed to detect editing or modifications [40]. Many machine learning based methods are employed to approach the problem in recent years [5, 41, 42]. The dataset that we are using in this work was originally proposed in [42]. It contains 300 length RNA sequences with 125 positive and 119 negative sequences.

#### 2.1.4 DNA Binding Proteins

DNA binding proteins bind to specific regions of DNA and affects the gene regulation. In this paper we have used a very well used benchamrk dataset for DNA binding proteins with 525 positive and 550 negative samples. This dataset was originally proposed in [43] and has been used extensively in the literature [7, 6, 44, 43, 45].

### 2.2 Feature Representation

After the data collection, the most important step in machine learning based methods is to convert the problem instances to vector representation. Generally, the feature vector is a collection of properties.

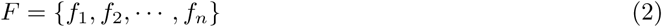

Different feature representation techniques have been used in the literature that includes: structural information [8], evolution properties [22, 11],etc. However, in recent works it has been shown that sequence based features though very easy and simple to generate are most effective if selected or designed properly [15, 44]. Moreover, our main objective in this work was to provide a generic framework for all three types of the sequences and to reduce the complexity in the feature generation step. That is the reason that we have selected to use sequence based features only. However, the framework still supports other features based derived or secondary properties and usable wherever its necessary and useful.

For the sake of simplicity in the experiments, we have selected similar group of features for all three type of sequences: Monomer composition, di-mer composiotion, trimer compostion, 1-gapped di-mono composition and 1-gapped mono-di compositions. However, based on the alphabet size the number of features extracted is different. We have used PyFeat tool [15] for feature extraction. Considering no overlaps, these features are then divided intro three groups. The details of the features are given in Table 2 and Table 3.

**Table 2:** Details of feature subspacing for protein dataset.

| Feature Subspace | Feature Type | No. of features |
| --- | --- | --- |
| $F_1$ | MonoMer Composition | 20 |
|  | DiMer Composition | 400 |
|  | TriMer Composition | 8000 |
| $F_2$ | 1-Gapped Di-Mono Composition | 8000 |
| $F_3$ | 1-Gapped Mono-Di Composition | 8000 |

**Table 3:** Details of feature subspacing for DNA and RNA dataset.

| Feature Subspace | Feature Type | No. of features |
| --- | --- | --- |
| $F_1$ | MonoMer Composition | 4 |
|  | DiMer Composition | 16 |
|  | TriMer Composition | 64 |
| $F_2$ | 1-Gapped Di-Mono Composition | 64 |
| $F_3$ | 1-Gapped Mono-Di Composition | 64 |

### 2.3 *SubFeat* Algorithm

The pseudo-code of *SubFeat* algorithm is given in Algorithm 1. It follows the same procedure as described in Figure 1. However, given a set of instances in the training set, *X* and the labels associated with them *y*, the algorithm first extract the feature set, *F*. From, *F*, next it populates a feature subspace set, 𝕏_*s*_. This set contains all the subspaces and this is controlled by two parameters, *n*_*p*_ denoting the number of partitions in the feature space and *overlap* is a boolean indicating whether there will be overlaps among the subspaces or not. In practice, *n*_*p*_ and *overlap* could be hyper-parameters and needs to be trained based on a specific problem in concern. After that, iteratively the hypothesis set, 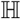 and associated weights, 𝕎 are learned based on the classifier type selected.

For prediction, the hypothesis set, 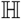 and weights set 𝕎 are used to ensemble the predictions of the individual base classifiers in a weighted majority fashion. The parameter *mix* allows the mix of the models selected.

### 2.4 Performance Evaluation

There are two important aspects of the performance evaluation: test sampling and metrics. In this paper, we have used 10-fold cross validation for the sampling of the datasets. The dataset is divided into 10 different balanced subsets retaining the balance ratio and then in each iteration 1 subset is used as test and the rest are taken as train set. These process in continued 10 times. However, to tackle the randomness effect, 10 runs were performed and average of them are reported only.

#### Algorithm 1: *SubFeat* (*X, y, n*_*p*_ = 3, *overlap* = *false*)

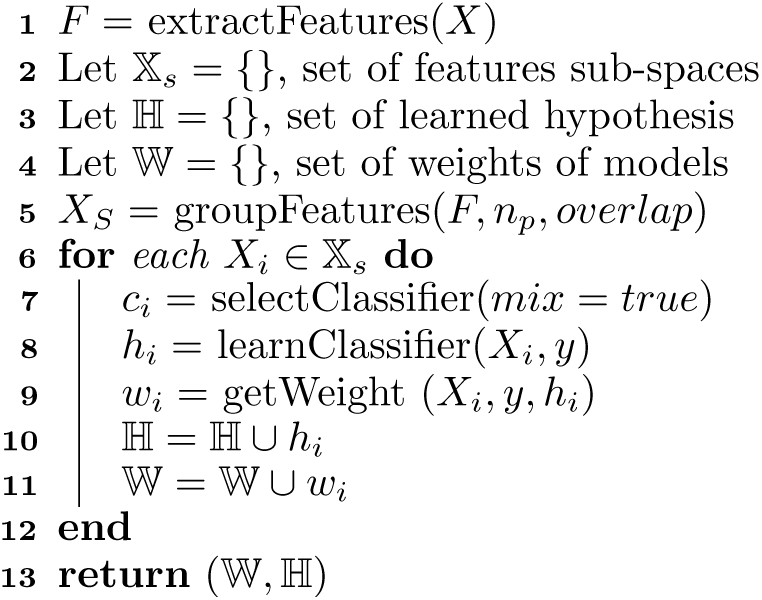

We have used several evaluation metrics: Accuracy (Acc), Precision, F1 Score, MCC, Sensitivity (Sn), Specificity and Area under curve (AUC). They are presented here in brief. Please note that, in the following equations TP, TN, FP and FN represents true positive, true negative, false positive and false negative. True positive means positive instances that were correctly classified by the classifier. True negative means negative instances that were correctly classified by the classifier. Similarly false positive and false negative means negative instances that are incorrectly classified as positive by the classifier and positive instances that are incorrectly classified as negative by the classifier.

1. **Accuracy (Acc)** gives a percentage result of correctly classified instances in between total number of instances.

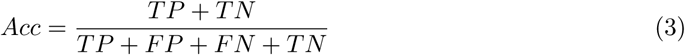
2. **Sensitivity (Sn)** gives a percentage result of correctly classified positive instances in between total number of positive instances.

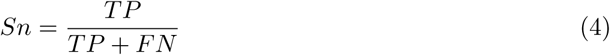
3. **Specificity (Spc)** gives a percentage result of correctly classified negative instances in between total number of negative instances.

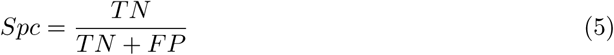
4. **Matthew’s Correlation Coefficient (MCC)** returns value between +1 to −1. The 0 represent a random classifier. The more the value is closer to +1, the better the classifier, similarly values towards −1 represent bad classifier.

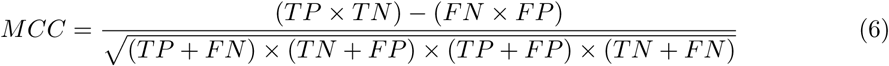
5. **F1 score** is the weighted average of precision and Recall. F1-score works with both false positive and false negative. Especially in the term of an uneven class distribution, this metric is usually more useful than accuracy.

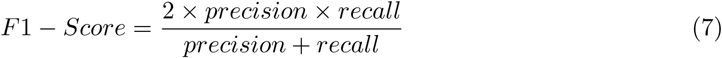 **Precision** gives a result of correctly classified positive instances in between total number of positive instances.

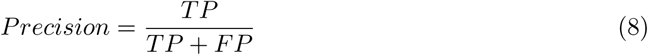 Recall is same as sensitivity and it is the ratio of correctly predicted true positive and false positive (all positive observations). It works on binary classification.

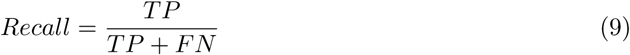
6. **Area under the Receiver Operating Characteristic curve** (AUC) is a performance measurement for classification problems at various thresholds. AUC is the measure or degree of separability while ROC represents a probability curve.

## 3 Results and Discussion

All experiments done in this paper are implemented in Python Language and using scikit learn library [46]. All experiments were run 10 times and the average of the results are reported. In all the tables bold faced values means the best values.

### 3.1 Classification Algorithms

In this section, we briefly describe the single based classifiers and the ensembles that were used for the experiments and for performance comparisons. Four single classifiers were used: Support Vector machines (SVM), Naive Bayes (NB), Decision Tree (DT) and Logistic Regression (LR). Support Vector Machine (SVM) [47] selects vectors that can represent the decision boundary best to separate the different classes. In our experiments, we have used a linear kernel based SVM. Logistic Regression (LR) [48] divides the sample space using linear hyper-planes. We use L2 regularization and regularization parameter set to 1.0 for the experiments with iterations to learn the parameters to 100. Decision Tree [49] is based on selecting features based on a measurement that can discriminate the instances best according to a criteria. We used*gini index* as the selection criteria and min samples to split was set 2. Gaussian Naive Bayesian (NB) [50] is a supervised learning based on probabilities of the features given the class labels and their likelihoods.

In addition to these single classifiers we have used three ensemble algorithms for experiments: Ad-aBoost, Random Forest and Ensemble Voting. Each of these algorithms are state-of-the-art ensemble methods that are used in the bioinformatics domain and as well as in other areas [27, 28].

### 3.2 Experimental Results

We present the results obtained by running experiments on four of the datasets. Table 4, Table 5, Table 6 and Table 7 shows the result of using single classification, feature subspacing ensemble classification and different ensemble classifier like random forest, adaboost and ensemble voting algorithms on Recombination Hotspots, *σ*^70^ promoters, RNA editing and DNA binding proteins problem respectively.

**Table 4:** Experimental Result on Recombination Hotspot Prediction Dataset.

| Algorithm | Precision | F1 | Acc | MCC | Sn | Spc | AUC |
| --- | --- | --- | --- | --- | --- | --- | --- |
| Result of single classifier algorithms |  |  |  |  |  |  |  |
| SVM | 0.8794 | 0.8016 | 0.7826 | 0.5633 | 0.7950 | **0.7673** | 0.8667 |
| NB | 0.7923 | 0.5975 | 0.6525 | 0.3636 | **0.8065** | 0.5794 | 0.7969 |
| LR | **0.8839** | **0.8034** | **0.7854** | **0.5687** | 0.8018 | 0.7658 | **0.8687** |
| DT | 0.7070 | 0.7547 | 0.7339 | 0.4659 | 0.7574 | 0.7061 | 0.7321 |
| Result of using different ensemble classifiers |  |  |  |  |  |  |  |
| Random Forest | **0.8913** | **0.8322** | **0.8120** | **0.6225** | 0.8098 | **0.8150** | **0.8874** |
| Adaboost | 0.8589 | 0.7982 | 0.7760 | 0.5492 | 0.7827 | 0.7672 | 0.8497 |
| Ensemble Voting | 0.8794 | 0.7754 | 0.7699 | 0.5471 | **0.8244** | 0.7186 | 0.8654 |
| Result of feature subsampling ensemble classification |  |  |  |  |  |  |  |
| SVM+SVM+SVM | **0.9724** | 0.8760 | 0.8464 | 0.7158 | 0.7835 | **0.9865** | **0.9708** |
| NB+Nb+Nb | 0.9681 | **0.9078** | **0.8946** | **0.7911** | **0.8674** | 0.9351 | 0.9647 |
| LR+LR+LR | 0.9697 | 0.8731 | 0.8420 | 0.7079 | 0.7787 | 0.9856 | 0.9706 |
| DT+DT+DT | 0.8562 | 0.8440 | 0.8297 | 0.6584 | 0.8420 | 0.8154 | 0.8771 |
| SVM+Nb+LR | 0.9505 | 0.8871 | 0.8632 | 0.7420 | 0.8072 | 0.9739 | 0.9423 |
| Nb+LR+SVM | 0.9498 | 0.8907 | 0.8676 | 0.7502 | 0.8113 | 0.9780 | 0.9441 |
| LR+SVM+Nb | 0.9471 | 0.8925 | 0.8697 | 0.7548 | 0.8128 | 0.9813 | 0.9421 |
| DT+SVM+DT | 0.9194 | 0.8689 | 0.8483 | 0.6980 | 0.8227 | 0.8884 | 0.9079 |
| SVM+DT+DT | 0.9148 | 0.8684 | 0.8481 | 0.6976 | 0.8233 | 0.8871 | 0.9065 |
| LR+LR+DT | 0.9199 | 0.8852 | 0.8609 | 0.7366 | 0.8060 | 0.9689 | 0.9047 |
| SVM+LR+DT | 0.9182 | 0.8824 | 0.8582 | 0.7286 | 0.8070 | 0.9562 | 0.9032 |
| SVM+Nb+DT | 0.9382 | 0.8916 | 0.8731 | 0.7513 | 0.8355 | 0.9354 | 0.9313 |

**Table 5:** Experimental Result on *σ*70 promoters dataset.

| Algorithm | Precision | F1 | Acc | MCC | Sn | Spc | AUC |
| --- | --- | --- | --- | --- | --- | --- | --- |
| Result of single classifier algorithms |  |  |  |  |  |  |  |
| SVM | 0.8924 | 0.8238 | 0.7647 | 0.4721 | 0.8070 | 0.6742 | 0.8229 |
| NB | 0.8883 | 0.7936 | 0.7473 | **0.4790** | **0.8501** | 0.6093 | 0.818 |
| LR | **0.8978** | **0.8262** | **0.7655** | 0.4696 | 0.8013 | **0.6835** | **0.8286** |
| DT | 0.7377 | 0.7604 | 0.6881 | 0.3142 | 0.7638 | 0.5490 | 0.6574 |
| Result of using different ensemble classifiers |  |  |  |  |  |  |  |
| Random Forest | **0.9024** | **0.8331** | **0.7735** | 0.4862 | 0.8036 | **0.7020** | **0.8368** |
| Adaboost | 0.8848 | 0.8106 | 0.7490 | 0.4399 | 0.7997 | 0.6452 | 0.8084 |
| Ensemble Voting | 0.8967 | 0.8188 | 0.7652 | **0.4865** | **0.8255** | 0.6563 | 0.8243 |
| Result of feature subsampling ensemble classification |  |  |  |  |  |  |  |
| SVM+SVM+SVM | 0.9589 | **0.8860** | 0.8098 | 0.5664 | 0.8007 | **0.8408** | **0.9232** |
| NB+Nb+Nb | 0.9513 | 0.8556 | **0.8203** | **0.6255** | **0.9008** | 0.7038 | 0.9084 |
| LR+LR+LR | **0.9598** | 0.8552 | 0.7886 | 0.5170 | 0.7745 | 0.8470 | 0.9222 |
| DT+DT+DT | 0.8261 | 0.8227 | 0.7605 | 0.4577 | 0.7975 | 0.6758 | 0.7786 |
| SVM+Nb+LR | 0.9442 | 0.8680 | 0.8175 | 0.5853 | 0.8233 | 0.8020 | 0.8969 |
| Nb+LR+SVM | 0.9446 | 0.8663 | 0.8153 | 0.5796 | 0.8225 | 0.7962 | 0.8964 |
| LR+SVM+Nb | 0.9443 | 0.8670 | 0.8166 | 0.5836 | 0.8240 | 0.7970 | 0.8964 |
| DT+SVM+DT | 0.9007 | 0.8406 | 0.7791 | 0.4935 | 0.7958 | 0.7336 | 0.8275 |
| SVM+DT+DT | 0.9021 | 0.8447 | 0.7857 | 0.5101 | 0.8023 | 0.7413 | 0.8320 |
| LR+LR+DT | 0.9178 | 0.8508 | 0.7862 | 0.5079 | 0.7829 | 0.7980 | 0.8414 |
| SVM+LR+DT | 0.9222 | 0.8563 | 0.7960 | 0.5326 | 0.7939 | 0.8031 | 0.8545 |
| SVM+Nb+DT | 0.9297 | 0.8602 | 0.8119 | 0.5765 | 0.8367 | 0.7566 | 0.8736 |

**Table 6:** Experimental Result on RNA editing dataset.

| Algorithm | Precision | F1 | Acc | MCC | Sn | Spc | AUC |
| --- | --- | --- | --- | --- | --- | --- | --- |
| Result of single classifier algorithms |  |  |  |  |  |  |  |
| SVM | 0.8788 | 0.7809 | 0.7918 | 0.5894 | 0.8019 | 0.7832 | 0.860 |
| NB | 0.8263 | 0.7359 | 0.7546 | 0.5165 | 0.7706 | 0.7407 | 0.7990 |
| LR | **0.9021** | **0.8041** | **0.8128** | **0.6342** | **0.8182** | **0.8088** | **0.8823** |
| DT | 0.6627 | 0.7087 | 0.7224 | 0.4535 | 0.7187 | 0.7256 | 0.7219 |
| Result of using different ensemble classifiers |  |  |  |  |  |  |  |
| Random Forest | 0.8801 | 0.7379 | 0.7765 | 0.5724 | **0.8500** | 0.7315 | 0.8483 |
| Adaboost | 0.8153 | 0.7217 | 0.7409 | 0.4901 | 0.7476 | 0.7357 | 0.7887 |
| Ensemble Voting | **0.9009** | **0.7779** | **0.7965** | **0.6048** | 0.8184 | **0.7794** | **0.8775** |
| Result of feature subsampling ensemble classification |  |  |  |  |  |  |  |
| SVM+SVM+SVM | **0.9315** | 0.8007 | 0.8310 | 0.6833 | **0.9225** | 0.7764 | 0.9137 |
| NB+NB+NB | 0.9155 | **0.8386** | **0.8500** | **0.7065** | 0.8694 | **0.8339** | 0.9059 |
| LR+LR+LR | 0.9302 | 0.8048 | 0.8276 | 0.6680 | 0.8860 | 0.7882 | **0.9144** |
| DT+DT+DT | 0.8251 | 0.8070 | 0.8106 | 0.6280 | 0.7993 | 0.8223 | 0.8619 |
| SVM+NB+LR | 0.8932 | 0.8070 | 0.8283 | 0.6692 | 0.8813 | 0.7904 | 0.88024 |
| NB+LR+SVM | 0.9012 | 0.7932 | 0.8219 | 0.6598 | 0.8892 | 0.7780 | 0.8896 |
| LR+SVM+NB | 0.9002 | 0.8060 | 0.8288 | 0.6704 | 0.8860 | 0.7890 | 0.8831 |
| DT+SVM+DT | 0.8993 | 0.8263 | 0.8382 | 0.6843 | 0.8647 | 0.8176 | 0.8876 |
| SVM+DT+DT | 0.8900 | 0.8106 | 0.8243 | 0.6553 | 0.8448 | 0.8083 | 0.8796 |
| LR+LR+DT | 0.8974 | 0.8116 | 0.8293 | 0.6686 | 0.8677 | 0.8011 | 0.8840 |
| SVM+LR+DT | 0.8779 | 0.7837 | 0.8097 | 0.6351 | 0.8659 | 0.7716 | 0.8584 |
| SVM+NB+DT | 0.8846 | 0.7999 | 0.8179 | 0.6659 | 0.8533 | 0.7918 | 0.8686 |

**Table 7:** Experimental Result on DNA binding proteins dataset.

| Algorithm | Precision | F1 | Acc | MCC | Sn | Spc | AUC |
| --- | --- | --- | --- | --- | --- | --- | --- |
| Result of single classifier algorithms |  |  |  |  |  |  |  |
| SVM | 0.7925 | 0.6986 | 0.7108 | 0.4279 | 0.7472 | 0.6812 | 0.7849 |
| NB | 0.5512 | 0.6643 | 0.5754 | 0.1623 | 0.5577 | 0.6297 | 0.5708 |
| LR | **0.8129** | **0.7303** | **0.7333** | **0.4696** | **0.7555** | **0.7130** | **0.7995** |
| DT | 0.5864 | 0.6273 | 0.6189 | 0.2387 | 0.6272 | 0.6103 | 0.6187 |
| Result of using different ensemble classifiers |  |  |  |  |  |  |  |
| Random Forest | **0.7821** | 0.7072 | **0.7000** | **0.4009** | **0.7058** | 0.6940 | **0.7769** |
| Adaboost | 0.7145 | 0.6760 | 0.6673 | 0.3358 | 0.6734 | 0.6611 | 0.7190 |
| Ensemble Voting | 0.7768 | **0.7181** | 0.6922 | 0.3879 | 0.6753 | **0.7160** | 0.7583 |
| Result of feature subsampling ensemble classification |  |  |  |  |  |  |  |
| SVM+SVM+SVM | **0.9051** | 0.7741 | 0.7227 | 0.4833 | 0.6641 | **0.8697** | **0.9004** |
| NB+Nb+Nb | 0.6075 | 0.6908 | 0.5990 | 0.2256 | 0.5704 | 0.7042 | 0.6440 |
| LR+LR+LR | 0.8788 | **0.8128** | **0.7903** | **0.5905** | **0.7488** | 0.8542 | 0.8822 |
| DT+DT+DT | 0.6617 | 0.6694 | 0.6634 | 0.3276 | 0.6728 | 0.6538 | 0.6987 |
| SVM+Nb+LR | 0.7923 | 0.7620 | 0.7105 | 0.4524 | 0.6578 | 0.8363 | 0.7618 |
| Nb+LR+SVM | 0.7914 | 0.7602 | 0.7071 | 0.4465 | 0.6543 | 0.8357 | 0.7644 |
| LR+SVM+Nb | 0.7900 | 0.7568 | 0.7055 | 0.4406 | 0.6550 | 0.8234 | 0.7615 |
| DT+SVM+DT | 0.7791 | 0.7288 | 0.7041 | 0.4117 | 0.6861 | 0.7291 | 0.7647 |
| SVM+DT+DT | 0.7810 | 0.7365 | 0.7120 | 0.4277 | 0.6918 | 0.7403 | 0.7714 |
| LR+LR+DT | 0.7914 | 0.7770 | 0.7538 | 0.5135 | 0.7241 | 0.7976 | 0.7744 |
| SVM+LR+DT | 0.7934 | 0.7715 | 0.7427 | 0.4944 | 0.7071 | 0.7993 | 0.7734 |
| SVM+Nb+DT | 0.7704 | 0.7258 | 0.6735 | 0.3639 | 0.6362 | 0.7525 | 0.7477 |

#### 3.2.1 Recombination Hotspot Prediction

For the Recombination hotspot prediction dataset, the results are presented in Table 4. The first part of the table shows that among the single classifiers, Logistic Regression performs significantly close. Since SVM and LR both are using linear decision boundaries, there performance very close to each other. However, when we turn to ensembles, we could notice Random Forest algorithm performs significantly better compared to other methods. In the lower part of the table, we present the results obtained by *SubFeat* using different combinations of single base classifiers. Note that, for this paper we have used only three base classifiers. Performance of all decision tree combinations is somewhat poor compared to others. Among all these combinations it appears that Naive Bayes and SVM combinations are working best. Here we can conclude that the mix of the base classifiers are not working well as compared to the combination of using the same base classifiers. Also note that these results by a good margin better over the results obtained by the ensemble methods.

Figure 2 shows area under receiver operating characteristic curves analysis for the recombination hotspot dataset. In this figure, we also put the standard deviations among all the runs. We could notice that here too, the proposed method shows higher performance and over the different thresholds its performance is superior to the other methods, single or ensemble. The strong performance of the proposed method, *SubFeat* in terms of AUC provides evidence on the robustness of the method.

**Figure 2:**
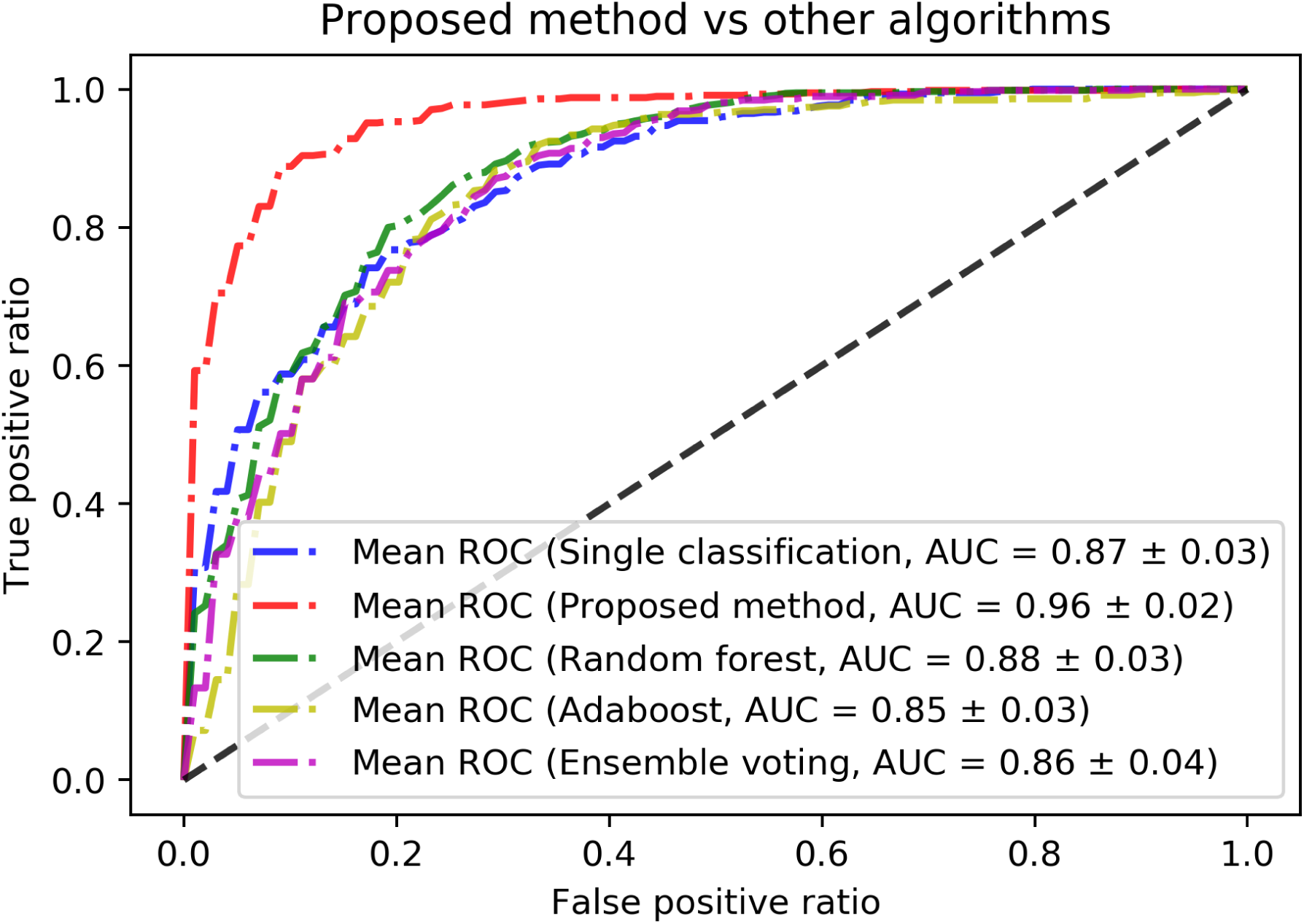
ROC analysis for recombination hotspot problem dataset.

#### 3.2.2 *σ*70 Promoters Prediction

Table 5 presents the results of our experiments on the *σ*^70^ promoters prediction problem. Here too we have presented the results in three parts: single, ensembles and *SubFeat* and its variations. From the results obtained in the single classifier experiments, we note that logistic regression outperforms the other methods. However, once again the performance of SVM is very close to logistic regression which is expected. In the ensemble part the results are improved compared to the single classifier results. Here, we could notice that Random FOest outperforms the rest of the methods. Moving to the third part of the table, we find the results of the different combinations of the single classifiers within the *SubFeat* framework. Similar to the results on the recombination hotspot problem, here too we notice that the mix combination of the single classifiers are not working as compared to the ensemble created with same type of the classifier. The best performing combination was produced by Naive Bayes algorithm. SVM and logistic regression followed closely. Decision tree combinations performed poorly. Also note that this dataset was the largest among the datasets considered for this work.

The receiver operating characteristic analysis on the *σ*70 promoters prediction dataset are presented using a curve of false positive rate against true positive rate and shown in Figure 3. *SubFeat* method here outperforms the other methods with a good margin again. Note that the changes in the threshold on the x-axis of the curve does not change the true positive rates. For a balanced dataset chosen for the purpose, this is a strong indication of the superior performance of *SubFeat* over the other methods compared in this work.

**Figure 3:**
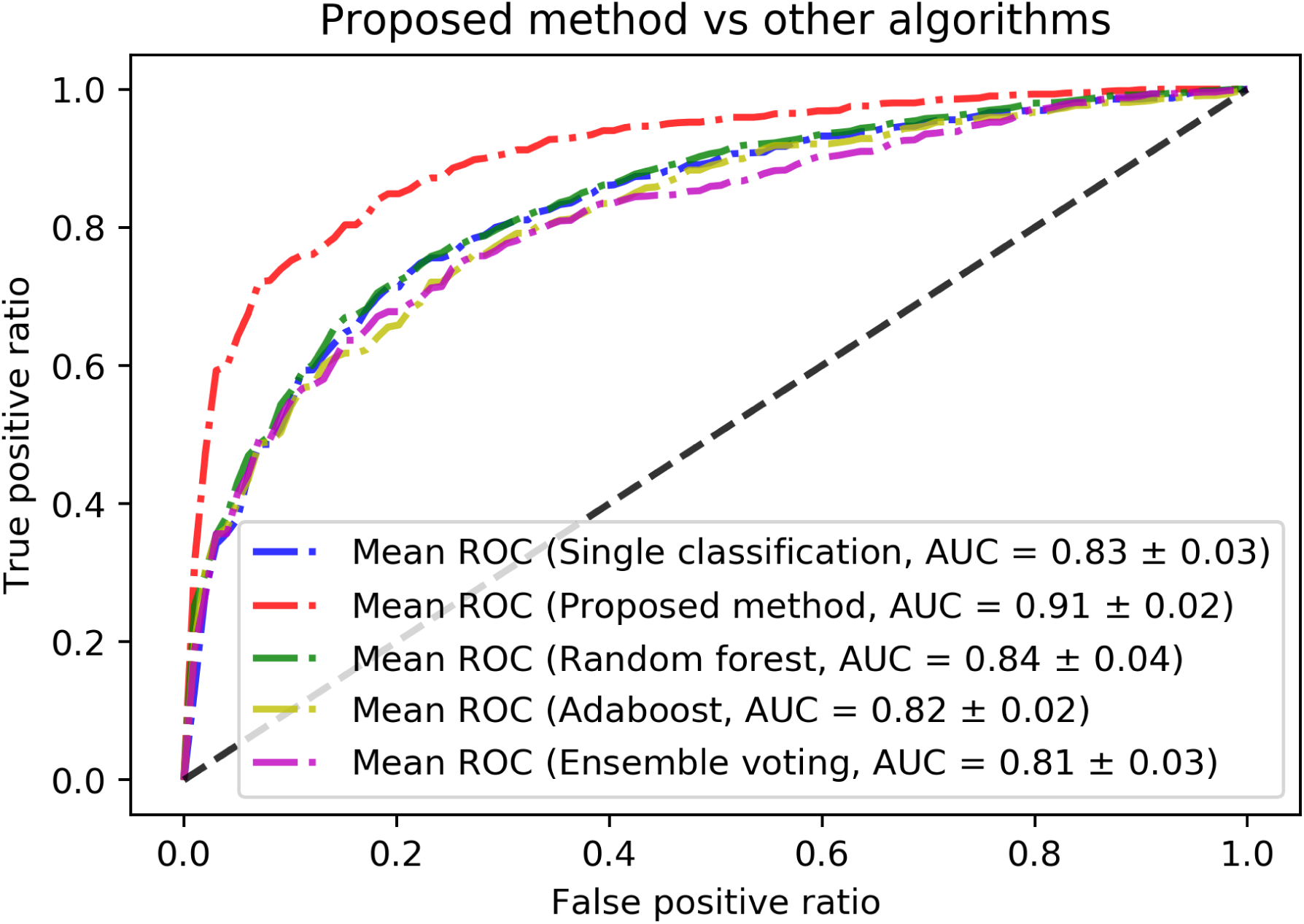
ROC analysis for *σ*70 promoters problem dataset.

#### 3.2.3 A-to-I RNA Editing Site Prediction

We present the experimental results on the A-to-I RNA editing sites prediction problem in Table 7. Note that, this is relatively smaller dataset compared to the other datasets. Here the performance of the single classifiers shown in the first part of the table are dominated by the logistic regression classifier in terms of all the performance metrics. Here, among the ensemble methods ensemble voting method performs significantly better compared to Random Forest or AdaBoost algorithms. However, *SubFeat* once again outperforms all these methods in terms of performance. This is clearly shown in the values reported in the lower part of the table. Here, we see that *SubFeat* follows the same trend as the previous datasets, that it the ensemble is working better when same classifier is chosen as base classifier. However, Naive Bayes is performing slightly better and SVM and logistic regression follows closely.

The ROC analysis for this dataset is shown in Figure 4. Note that, for this dataset though *SubFeat* is still superior in performance in terms of AUC values, the difference is not that high as compared to the other datasets. Here, single classifier is working better compared to other datasets.

**Figure 4:**
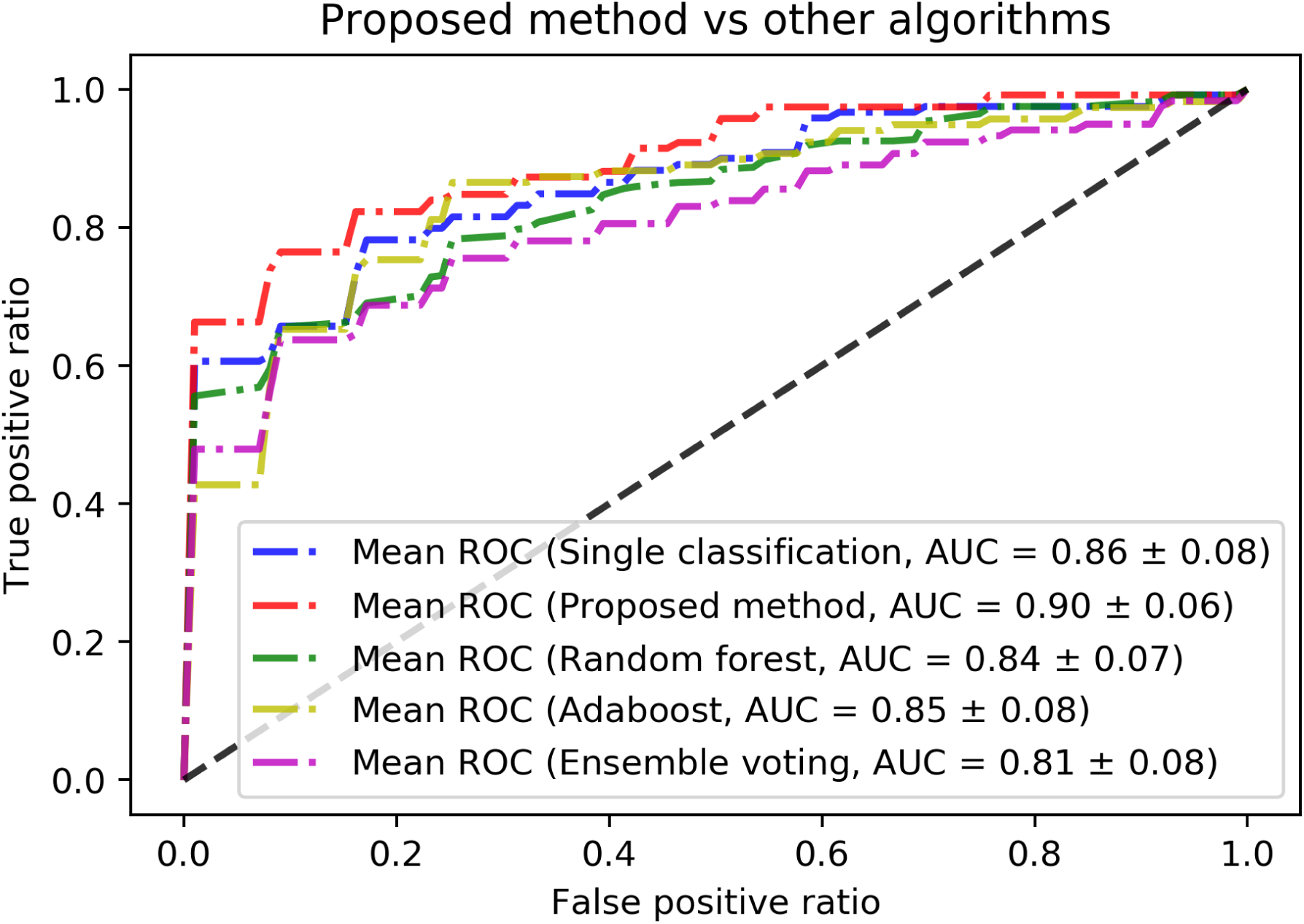
ROC analysis for RNA editing prediction problem dataset.

#### 3.2.4 DNA Binding Proteins Prediction

Experimental results on the DNA binding proteins prediction problem is reported in Table 7. We could note the similar trends for this dataset as well. Logistic regression performs best in the single classifier group. Similar to that performance combination of logistic regression classifier used in the *SubFeat* is best among all the classification algorithms. The performance of this combination is slightly weaker in terms of AUC comapred to the all SVM combination. This is due to the better precision values obtained by the SVM combination which is also reflected in the specificity values reported in the table. The ROC analysis is shown in FigureZ5 in more details. The plot shows the superior performance of *SubFeat* over all other methods.

**Figure 5:**
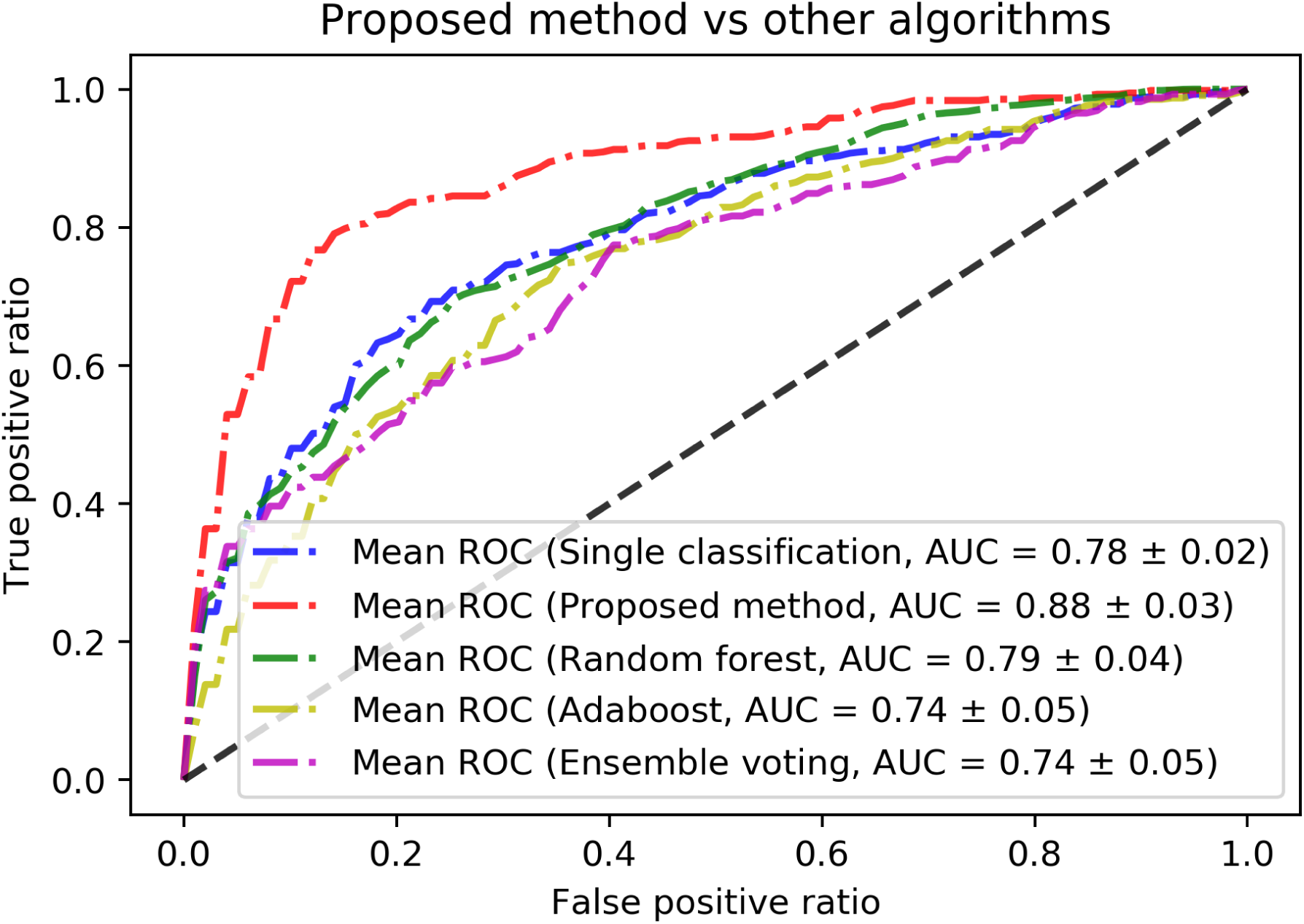
ROC analysis for DNA binding proteins prediction problem dataset.

### 3.3 Discussion

As a method, *SubFeat* shows better performance in all metrics compared to single and ensemble classifiers as found in the results and analysis shown in the previous section. That establishes the claim of the hypothesis of using an ensemble and dividing the feature space into subspaces. However, another subtle observation could be made from the results that using similar classifier as base classifier is achieving better results compared to the mix of the classifiers. This study was limited to four datasets and this remains still a question to be explored in details if the *mix* parameter can also bring good results. We believe that might be utilized as well. Two of the variables or parameters of the *SubFeat* framework is less explored in this paper. They are *n*_*p*_, number of partitions which is set to 3 in all the experiments and *overlap* which is kept false for all the experiments.

We believe answer to the performance largely depends on the feature space or the feature representation. In this work, we have limited to use only sequence based features. In problems like DNA binding protein prediction, we have noticed application of structural and evolutional features have been used successfully [7, 6]. In the cases of DNA and RNA sequences as well, the researchers have used many other types of feature representation technique. Note that the knowledge number of partitions for the feature space will obviously enhanced by selection of such techniques, as previously we have seen group based feature selection to be performing better in a wide range of problems [25, 8]. However, in those works, the idea of ensemble method was not explored. We kept the experimental setting simpler and thus not extended the feature space. We believe using a larger and enhanced feature space will improve the results.

Another parameter is the overlapping of the feature spaces. Though we have not reported the results, for these four datasets we have seen that the overlap parameters are not working well. We observed that sensitivity suffers of we accept overlap too much. Note that in a previous work [3], overlapping has been found effective for promoter prediction. The results presented in this paper are much superior compared to the ones reported in [3]. However, note that the objective of this paper is limited to show the effectiveness of the ensemble based on feature subspacing.

### 3.4 Python Package

We have made our method, *SubFeat* available as a Python based package. It is freely available for use from https://github.com/fazlulhaquejony/SubFeat. The package includes all the parameters that we have discussed and provided as option for the method. A simple to follow user guide is also provided on how to install and use the package along with example runs/experiments. We strongly believe that further exploration are possible for this package and it will be useful for the computational biologists working in the relevant fields.

## 4 Conclusion

In this paper, we have proposed a ensemble method where the full feature space was divided into subspaces. From the results we can conclude that the subspace method provide better prediction result compared to both the single classifiers and the best ensemble algorithms like AdaBoost, Random Forest, etc. We have tested the performance of the algorithm on a full space feature representation for protein, DNA and RNA sequences datasets. However, it is possible to improve our accuracy by using a different feature space and feature selection techniques. We have only tested our method on balanced binary classification biological datasets. We have tested using overlaps of the feature-spaces however the number of sub space is still a parameter to be tested comprehensively. Therefore, in future we plan to work with imbalance data, independent and large number of dataset. The simplicity of these method help to increase the accuracy of biological sequence datasets.

